# Sticky Situations: Bacterial Attachment Deciphered by Interferometry of Silicon Microstructures

**DOI:** 10.1101/793125

**Authors:** Heidi Leonard, Liran Holtzman, Yuri Haimov, Daniel Weizman, Sarel Halachmi, Yechezkel Kashi, Ester Segal

**Affiliations:** Department of Biotechnology and Food Engineering, Technion – Israel Institute of Technology, Haifa 3200003, Israel; Department of Urology, Bnai Zion Medical Center, Haifa 3104800, Israel; The Russell Berrie Nanotechnology Institute, Technion – Israel Institute of Technology, Haifa 3200003, Israel

**Keywords:** biointerfaces, bacteria, microstructures, sensors, photonics, biofilms

## Abstract

The peculiarities of surface-bound bacterial cells are often overshadowed by the study of planktonic cells in clinical microbiology. Thus, we employ phase-shift reflectometric interference spectroscopic measurements to observe the interactions between bacterial cells and abiotic, microstructured material surfaces in a label-free, real-time manner. Both material characteristics (i.e., substrate surface charge and wettability) and characteristics of the bacterial cells (i.e., motility, cell charge, biofilm formation, and physiology) drive bacteria to adhere to a particular surface. We conclude that the attachment of bacterial cells to a surface is determined by the culmination of numerous factors. When specific characteristics of the bacteria are met with factors of the surface, enhanced cell attachment and biofilm formation occur. Such knowledge can be exploited to predict antibiotic efficacy, biofilm development, enhance biosensor development, as well as prevent biofouling.

## 1. Introduction

Bacteria thrive in surface-associated communities, in which the colonized surface provides both protection and a stable anchor for the development of a dense cellular network.^[1]^ These populations of bacterial cells that adhere to a surface and to each other by a variety of secreted substances and appendages are often classified as biofilms. The latter can manifest themselves as medical nuisances and threats,^[2]^ such as antibiotic-resistant urinary tract infections or as part of natural ecosystems, such as the geochemical recycling of sediment.^[3,4]^ It has been reported that surface-associated bacterial networks are more resistant to environmental stresses, such as antibiotics, than their free-floating planktonic counterpart cells.^[5]^ Thus, elucidating and quantifying the attachment behavior of bacterial cells to their substrates and development of biofilms is crucial to mediating problems of deadly infections,^[6]^ understanding of fragile ecosystems, biofouling, and even the design of synthetic microbial populations.^[7–10]^

Although, traditionally, the behavior of microbes has been successfully studied in pure liquid cultures, new methods and materials for quantitatively examining bacterial attachment and colony formation on surfaces have recently emerged. Copious structurally hard and soft materials, specifically lamellar and nano-/micro- patterned materials, have been examined for their effect on bacterial colonization and adhesion;^[7,11–21]^ however, very few of these studies have employed the resulting intrinsic optical properties^[22]^ of the materials for real-time, quantitative evaluation of bacterial adhesion. Methods such as Raman spectroscopy,^[23]^ quartz crystal microbalances,^[24,25]^ and total internal reflectance fluorescence^[26,27]^ have been used to monitor bacterial adhesion and biofilm formation on surfaces, but employ substrates that can be difficult to chemically or structurally manipulate, while Raman spectroscopy only reveals the chemical signature of the bacteria,^[23]^ as opposed to a phenotypic quantification of adhered cells. Highly sensitive techniques, such as resonant mass sensors for cell characterization require the precise positioning of single cell on a cantilever tips,^[28]^ while supercritical-angle fluorescence requires immobilization of the specifically monitored cells.^[29]^ Furthermore, more traditional imaging techniques to elucidate bacterial behavior^[30]^ can be difficult to quantify cells, expensive, and destructive to the microbial networks.^[31]^

While many efforts have been focused on understanding mammalian cell interfaces with materials,^[32–35]^ in this work, we investigate bacterial cell responses to manipulated surfaces of silicon topologies by intrinsic phase-shift reflectometric interference spectroscopy measurements, termed PRISM.^[36,37]^ In this method, bacterial cells colonize on diffractive, patterned Si microstructures, while zero-order reflectance spectra are continuously collected during illumination with a broadband white light source.^[36–40]^ The resulting reflectance spectra exhibit optical interference fringes as the incident light is separated such that part of the beam is reflected by the top of the Si microstructures and the rest is reflected by the bottom of the Si microstructures. As bacteria occupy the empty spaces within the microstructures, the refractive index of the filling medium increases, resulting in increased values of optical path difference corresponding to *2nL*, in which *n* represents the refractive index of the filling medium and *L* refers to the height of the grating. Thus, bacterial adhesion to chemically and structurally modified substrates can be monitored in real time without the use of labels or sophisticated microscopes, allowing for the elucidation of long-disputed theories, such as the application of the Derjaguin–Landau–Verwey–Overbeek (DLVO) colloid stability theory to bacteria,^[41]^ and revealing tendencies of clinically relevant pathogenic bacteria. This extensive study of the abiotic-biotic interface incorporates a variety of bacteria types and shapes, ranging from common rod-shaped, laboratory strains of bacteria to pathogenic cocci of clinical isolates, on a variety of chemically functionalized microtopologies to highlight the paramount role of the material surface on the initiation of complex microbial communities.

## 2. Results & Discussion

### 2.1. PRISM monitors bacteria adhesion

PRISM assays to monitor bacterial adhesion are performed in transparent polycarbonate flow cells, each housing a Si photonic chip comprised of arrayed microstructures (**Figure 1**A), in the form of periodic pillars or pores (see Figure S1 for respective images of the gratings). For PRISM adhesion assays, the photonic chips are illuminated at a normal incident angle by a collimated broadband light source through the flow cell, while being exposed to various solutions at a flow rate of 20 μL min^−1^. Using frequency analysis of resulting reflectance spectra, the optical path difference between the two parts of the incident light beams is calculated as *Δ2nL* (%), or the percent change of *2nL*, over time (see Experimental for more details). Thus, as more bacterial cells collect within the microstructured grating, the *2nL* values increase as a function of the increasing refractive index (Figure 1B).

**Figure 1.**
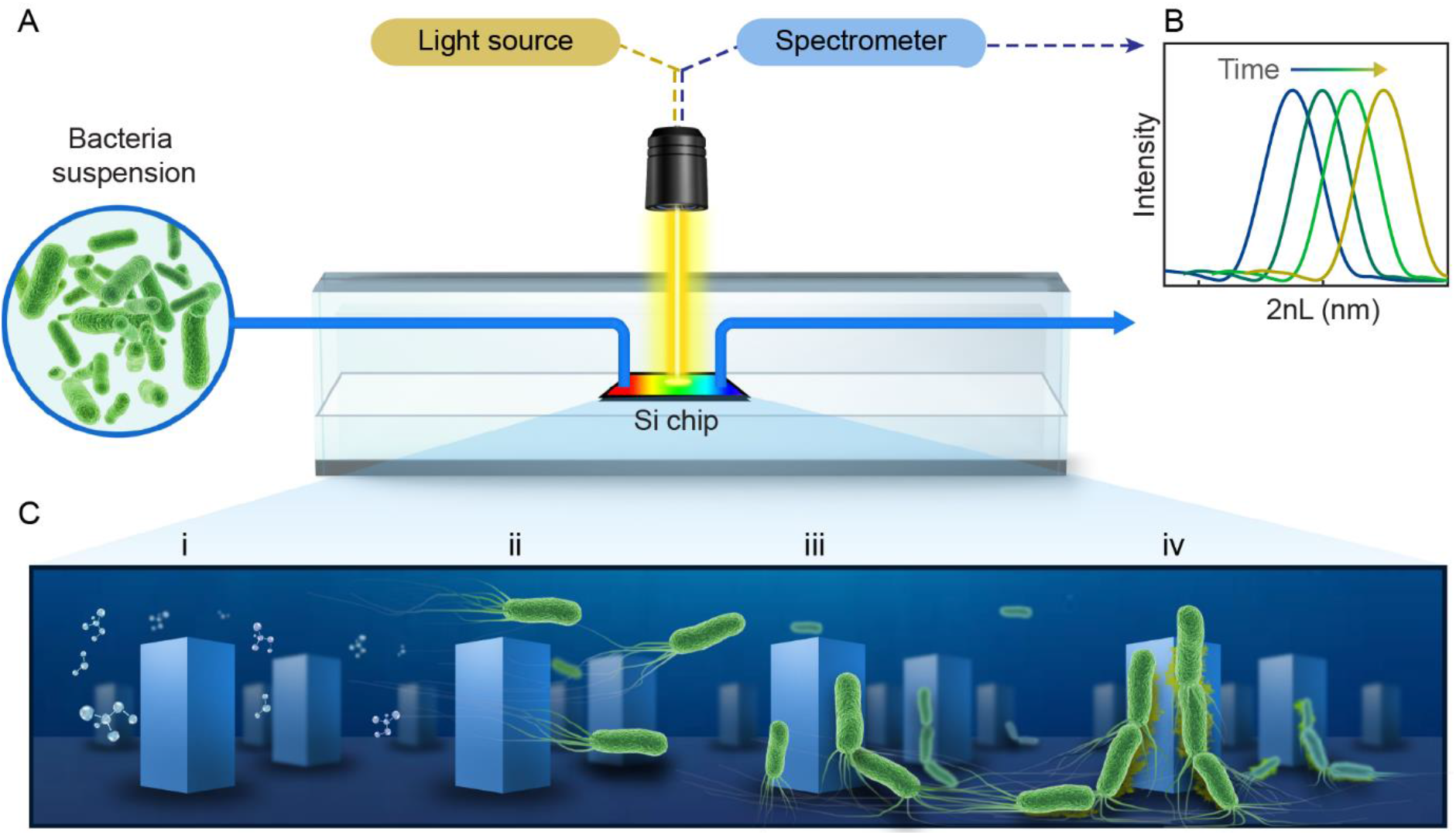
Schematic representation of PRISM for bacteria adhesion assays. (A) A bifurcated optic fiber, fitted with a collimating lens, illuminates a Si photonic chip while a bacteria suspension (composed in motility buffer, MB) is introduced in flow mode over the chip. A CCD detector continuously acquires zero-order reflectance spectra in which frequency analysis produces (B) the *2nL* values that increase over time as more bacteria adhere within the microstructured surface. (C) Four stages of bacteria attachment leading to biofilm formation are investigated, including (i) molecular preconditioning of the substrate microstructures during flow of sterile MB; (ii) transport of bacterial cells to the microstructured surface when the solution is changed to bacterial suspension; (iii) bacterial cell adhesion to the microstructures as the result of specific cell appendages and electrostatic charges; (iv) cohesion of cells to each other as they begin to form networks of biofilms.

By manipulating the topology and chemical functionality of the Si photonic chips, as well as examining the responses of a variety of bacteria species and strains, effects of different stages of bacteria attachment to the abiotic surfaces can be observed. Specifically, we focus on the attachment stages outlined in Figure 1C, which include the preconditioning of the microstructures with an aqueous solution of motility buffer (MB), followed by the arrival of cells to the topologies, their subsequent attachment to the surface, and the initial formation of a community as cells attach to each other.

### 2.2. Wetting and fluid flow precondition the substrate for bacteria

Molecular conditioning of a surface, the first step toward the formation of bacterial colonies, occurs before bacteria cells are present (Figure 1Ci). During this stage, the Si surface is introduced to an aqueous solution of MB and coated with ions from the surrounding medium - assuming sufficient wetting. The introduction of a liquid to a surface brings about two key players involved in bacteria cell attachment – the first being wettability,^[18]^ which is partially dependent on the structural topology of the surface^[42]^; and the second being surface charge, which is dependent on the surface chemistry.

Consequently, Si photonic chips containing one of two topologies (*i.e.*, pillars or pores) are utilized in order to observe the role of substrate wettability on bacteria colonization. The topologies are either kept as oxidized (termed as OX) surfaces, garnering a negative charge, or subject to amino-silanization (termed as AMINE), garnering a more positively charged surface in neutral-pH MB.^[43,44]^ Contact angle measurements conducted in MB on the OX Si topologies reveal that pillar structures exhibit a reduced contact angle value compared to planar substrates (56° vs. 80°, respectively), while pore topologies exhibit a greater contact angle value of 115°. The same trend is observed with AMINE-functionalized topologies as well, see **Figure 2A**. This suggests that the planar Si surfaces follow a Young wetting regime of a smooth surface, the wetting of pillars can be described by the Wenzel model, and the pore topologies follow the Cassie-Baxter model of wetting.^[45,46]^ While the Wenzel model suggests that liquid on a rough surface (*i.e.*, micropillars) can infiltrate entirely, the Cassie-Baxter model for pore topologies implies that air is trapped underneath the buffer within the pores, resulting in an air-liquid interface of micro-pockets of vapor, as schematically illustrated in Figure 2A.

**Figure 2.**
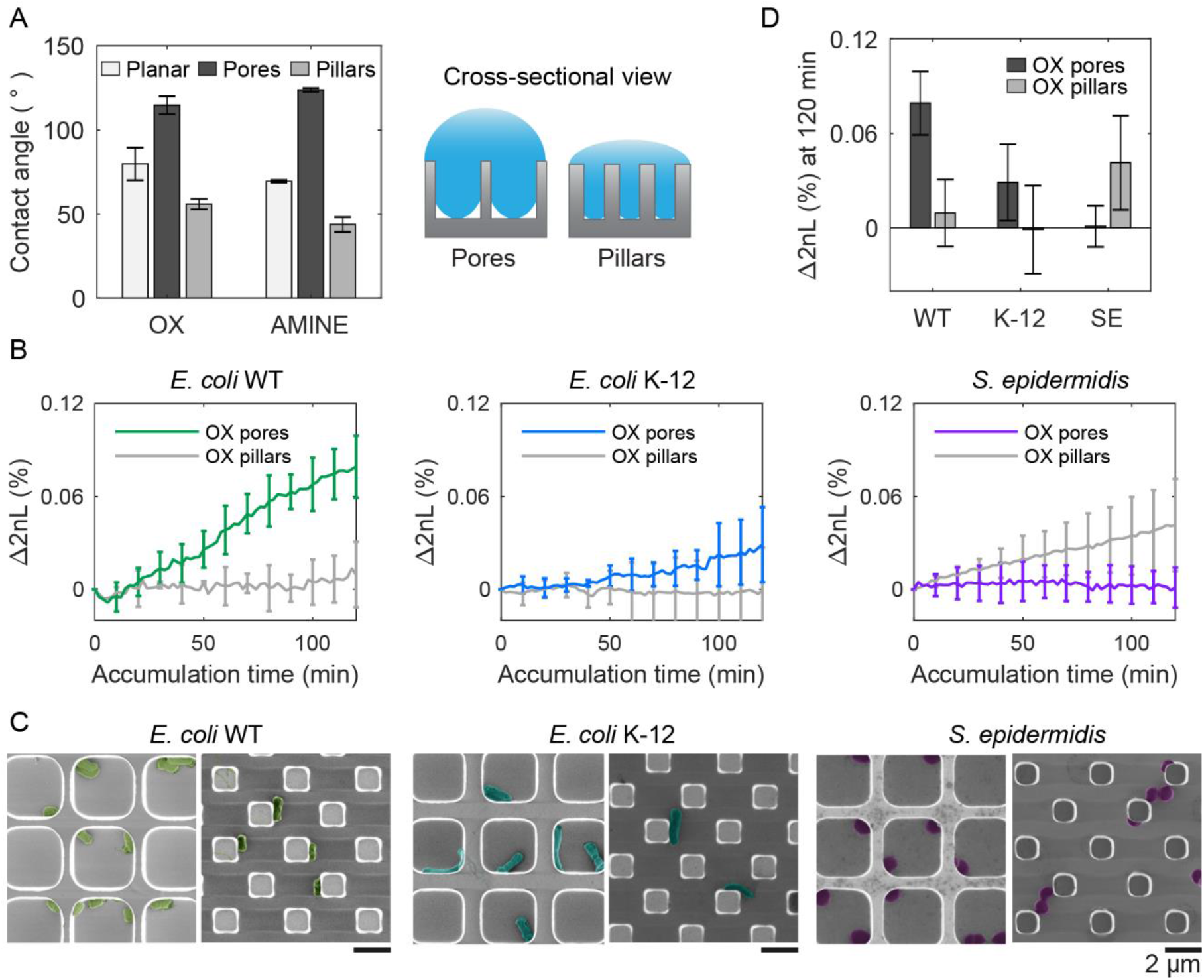
Different microtopologies promote varying degrees of wetting, in turn promoting differences in bacterial adhesion amongst strains. (A) Contact angle values (left panel) of MB measured after 200 s on oxidized (OX) and amine-functionalized (AMINE) pores, pillars, and planar Si substrates suggest that the buffer does not completely infiltrate the pore topologies, as illustrated (right panel). MB positions partially within the microtopologies, forming small vapor pockets in pore structures, characteristic of a Cassie-Baxter wetting regime. (B) PRISM assays of *E. coli* WT, *E. coli* K-12, and *S. epidermidis* adhesion to OX Si pores and Si pillars over time (n = 3). Measurements were collected every 2 minutes, but error bars are placed every 10 minutes for visual clarity. (C) False-colored SEM images of bacterial cells within the microstructured pores (left) and pillars (right) reveal attachment behavior to the oxidized surfaces for each of the different strains. Scale bars represents 2 μm. (D) Summary of the *Δ2nL* (%) values for each *E. coli* WT, *E. coli* K-12, and *S. epidermidis* (abbreviated SE) after 120 min of accumulation onto oxidized pores and pillars (n = 3).

During PRISM assays, Si chips are first exposed to sterile MB without bacteria in order to acquire a stable baseline signal. Thus, due to their intrinsic wetting properties, pillar and pore topologies already provide a different liquid environment before bacteria cells are introduced. As preliminary tests, the PRISM responses of three non-pathogenic bacteria strains, namely the laboratory mutant *Escherichia coli* (*E. coli*) K-12, clinically isolated *E. coli* (ATCC 25922, termed as WT), and *Staphylococcus epidermidis* (*S. epidermidis* ATCC 14990), were observed over time, with *Δ2nL* (%) corresponding to the accumulation of cells in the topologies (Figure 2B). By utilizing MB, as opposed to nutrient-rich growth medium, only events related to bacterial attachment are observed during the experiment, though cell motility is preserved.^[47,48]^

As complimentary analysis, scanning electron microscopy (SEM) images of the Si chips after 120 min of bacteria accumulation during PRISM assays (Figure 2C) reveal how individual *S. epidermidis* cells position themselves within the corners of the OX pores, while in the OX pillar topologies, *S. epidermidis* cells tend to cohere to other cells as opposed to the microstructures. Contrarily, rod-shaped *E. coli* cells position themselves alongside the OX pillars in straight lines, while in OX pores, they adhere either to the corners (as with *E. coli* WT) or in the middle of the pore (as with *E. coli* K-12). The microscopic menisci formed in the pore substrates underneath the MB may also explain why *S. epidermidis* cells tend to aggregate in the corners of the pores (Figure 2C), as it has been previously observed that certain types of bacteria prefer to congregate at air-liquid interfaces^[49]^ or perhaps this is where they accumulate the most according to Brownian motion.^[16]^ It is intriguing that not only are differences in adhesion noticeable between bacterial species, but also amongst bacterial strains, confirming that cell adhesion to a surface is a complex process.

Furthermore, Figure 2D summarizes PRISM adhesion assays for *E. coli* WT, *E. coli* K-12, and *S. epidermidis* accumulated after 120 min on OX pore and pillar substrates. According to these assays, both strains of *E. coli* (WT and K-12) exhibit a higher degree of accumulation within OX microstructured pores than to the OX pillar substrates, while *S. epidermidis* cells favor adhesion to OX pillar substrates over OX pore substrates. This confirms that even before the presence of bacteria, wetting of the pores and pillars provide different surrounding micro-environments that ultimately affect bacteria adhesion. It is also interesting to note that the *E. coli* WT exhibits more adhesion to oxidized pores than *E. coli* K-12, despite being of the same bacterial species. Thus, because wetting cannot be the only factor that affects bacterial adhesion, we proceed by investigating the role of cell motility and cellular phenotype in bacterial adhesion.

### 2.3. Bacterial mechanics guide cells to the surface

Noting the opposite adhesion preferences to pore and pillar substrates between species, differences in magnitude of attachment between species, and knowing that in order to adhere to a surface, bacterial cells must reach a close enough distance to attach, we investigated the role of bacteria motility. More specifically, we determine whether cells reach the microtopologies as part of active navigation guided by cell mechanics (e.g. flagellar motors and chemotactic responses) or if the cells passively fall into the interstitial space of the microstructures. By employing PRISM, genetically-modified *E. coli* mutants in the presence of OX substrates are monitored (**Figure 3**). Specifically, non-motile *E. coli* without flagella (*ΔflaA-flaH*, HCB137), motile *E. coli* without functional chemotactic receptors (*ΔcheA-cheZ*, HCB437), and *E. coli* with an incessant tumbling motion (*ΔcheZ*, HCB1394) are used as model bacterial strains for these studies.

**Figure 3.**
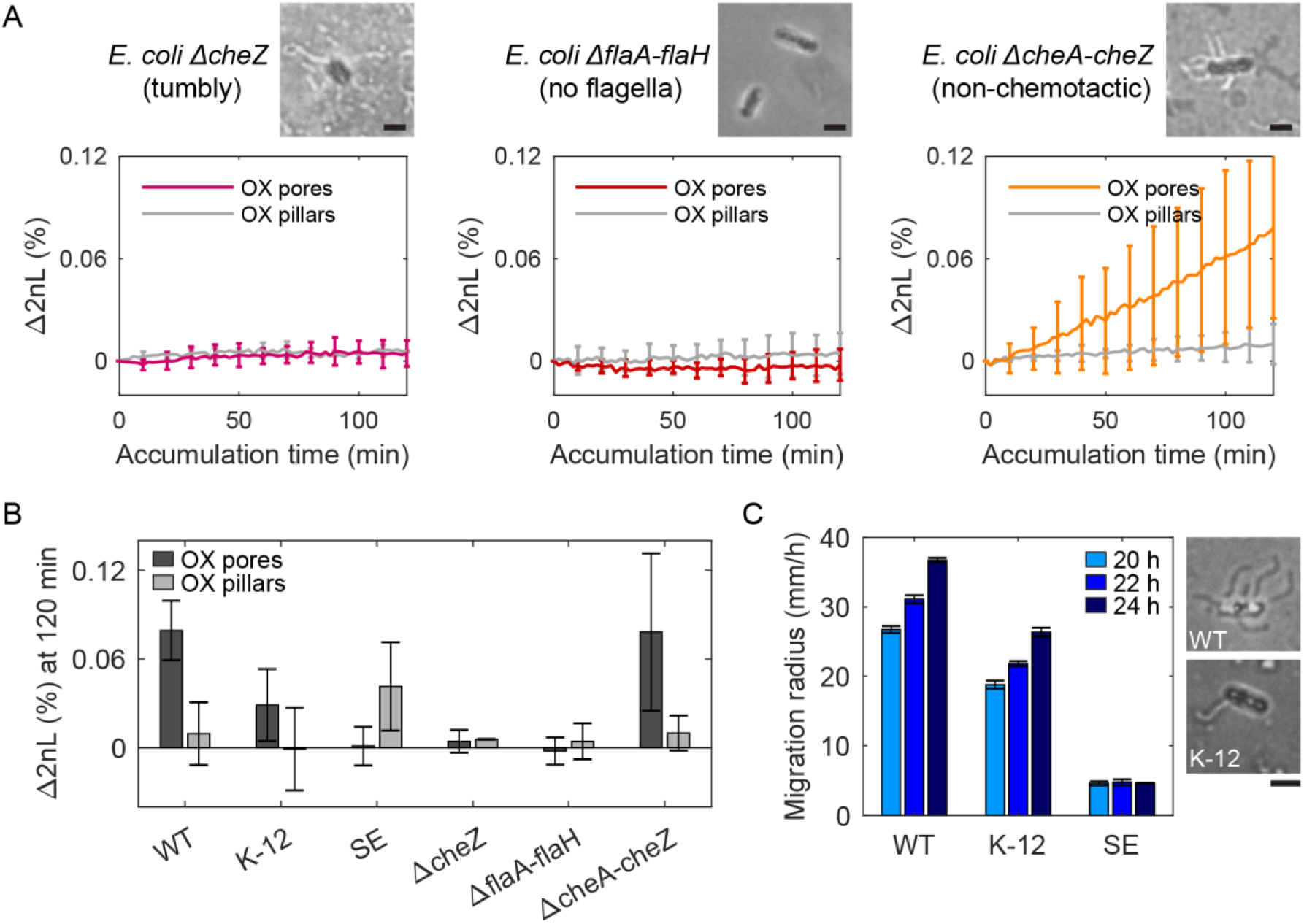
Motility is an underlying factor leading to bacterial adhesion within the microstructures. (A) PRISM of *E. coli* with a cheZ deletion causing excessive tumbling, *E. coli* without flagella, and *E. coli* with deleted chemotaxis receptors exhibit differences in adhesion behavior to oxidized (OX) pore and pillar substrates. Ryu flagellar staining reveals the presence or absence of flagella on the strains in optical microscope images. Scale bars represents 1 μm. (B) Summary of final *Δ2nL* (%) values reached after 120 min for the genetically modified *E. coli* mutants, *E. coli* WT, *E. coli* K-12, and *S. epidermidis.* (C) Semi-soft agar motility assays of *E. coli* WT, *E.* coli K-12, and *S. epidermidis* reveal that *E. coli* WT is significantly more motile than the K-12 strain (n = 4), while *S. epidermidis* is immotile. Optical microscope images after Ryu staining of *E. coli* WT and *E. coli* K-12 (right) reveal that the K-12 cells tend to have fewer flagella present. Scale bar represents 2 μm.

The least adhesive bacteria, tumbly *E. coli ΔcheZ*, did not adhere to either pillar or pore substrates as demonstrated by PRISM (Figure 3A), as they possibly do not have a long enough residence time to adhere to the surface. *E. coli ΔflaA-flaH* without flagella, exhibited similar behavior to *S. epidermidis* (see Figure 2B, left panel), which is also non-motile, with a slight preference towards OX pillar structures, but with lower levels of adhesion (Figure 3B). The significant difference in cells size between *E. coli* and *S. epidermidis*, may explain the lower degree of adhesion of the *E. coli ΔflaA-flaH* in pillars. We speculate the larger *E. coli* are unable to fit within the pillars as easily as the smaller *Staphylococci*. Contrarily, the non-chemotactic strain of *E. coli* behaved similar to the WT strain, confirming that the movement of bacteria into the microstructures during the PRISM assays is not directed by chemotaxis.

Furthermore, the motility of the *E. coli* WT, *E. coli* K-12, and *S. epidermidis* strains are measured in a semi-soft agar assay, demonstrating that *E. coli* WT swims faster than *E. coli* K-12, while *S. epidermidis* proves to be immotile (Figure 3C). This suggests that *S. epidermidis* passively falls into the microstructures as the result of Brownian motion and that gravitational force drives the cells to the bottom of an aqueous solution.^[50]^ On the contrary, both strains of *E. coli* actively accumulate in pore structures. However, *E. coli* WT exhibits a higher degree of accumulation, possibly due to the faster rate of swimming. Differences in swimming speeds between the two *E. coli* strains could possibly be explained by the varying number of flagella per cell as seen in optical microscope images (Figure 3C) after Ryu flagellar staining.^[51,52]^

Cell collisions with the microstructures are evidently crucial for attachment, though to an extent, the uncontrolled motility of bacteria may work against adhesion, as seen with *E. coli* ΔcheZ.^[53]^ In summary, while flagellar motors are not crucial for bacterial adhesion to the microstructured surface, they certainly assist in the process, particularly to pore topologies deficient in wetting and especially for larger sized cells that cannot fit within the confines of the pillars. These results also suggest that cells can passively diffuse between the pillar topology, but must actively swim into the pores to attach, unless congregating at the air-liquid interface of the micro-menisci.

### 2.4. Electrostatic forces promote cell adhesion at the surface

Once within the vicinity of the microstructures, bacterial adhesion is achieved by cellular appendages and secretions, but is also thought to be promoted by physiochemical forces.^[54]^ In particular, when bacteria reach a certain distance from the surface, they can then undergo reversible and irreversible attachment.^[50]^ Many studies try to draw parallels between the attraction of colloidal particles based on surface charge (described by DLVO theory) with bacteria cells adhering to abiotic surface.^[54]^ To test this association, we compare bacterial adhesion to positively and negatively charged substrates using PRISM (Figure 4). OX Si surfaces exhibit an overall negative charge due to the presence of hydroxyl groups, while AMINE surfaces exhibit an overall positive charge in pH-neutral MB (Figure 4A). The overall surface charge of each of the studied bacterial strains is also measured as a function of electrophoretic mobility by zeta potential measurements in MB (Figure 4B).

**Figure 4.**
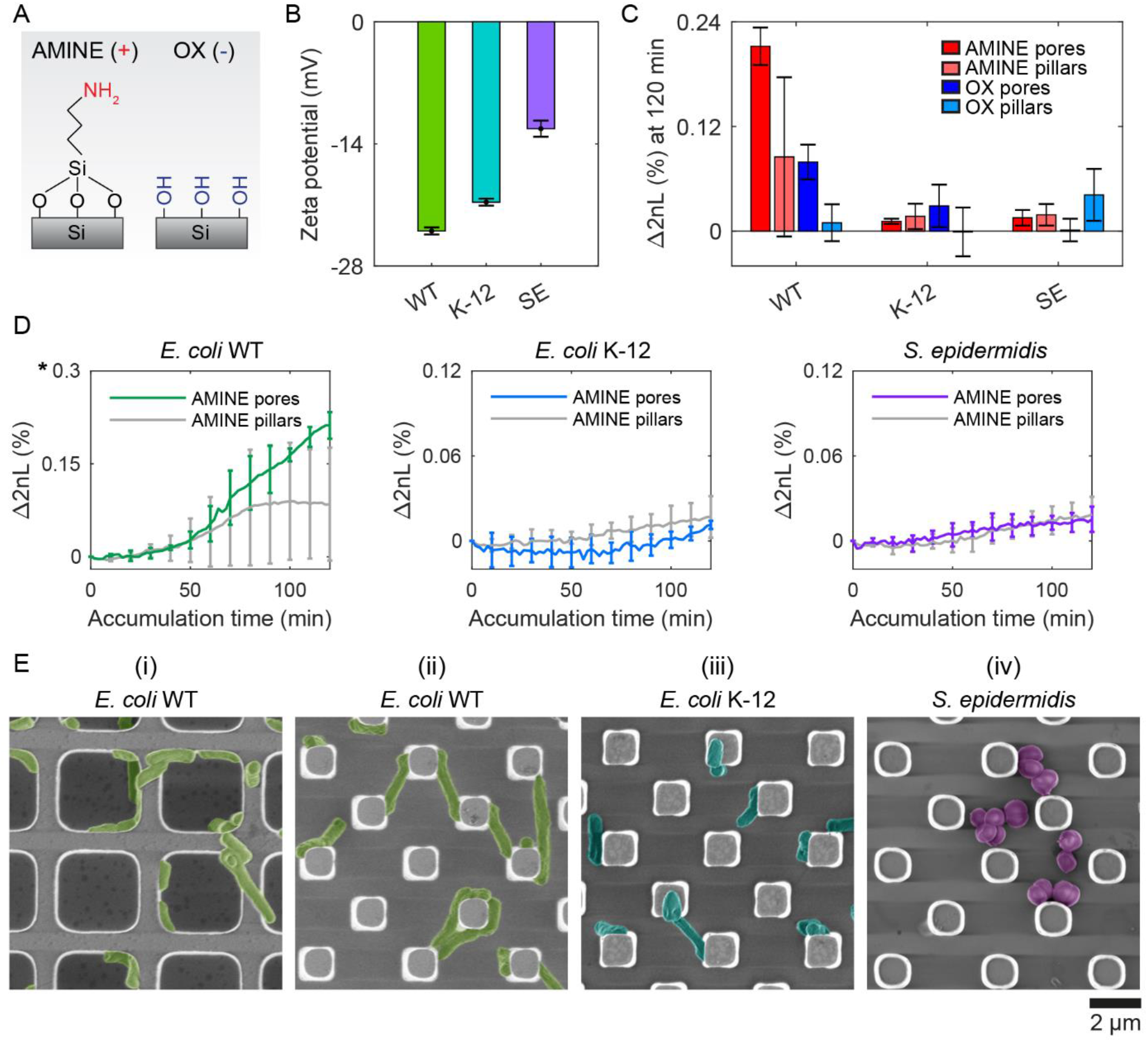
Substrate surface charge plays a role in bacterial adhesion. (A) AMINE Si surfaces yield a positive surface charge, while OX surfaces yield a negatively charged surface in motility buffer. (B) Zeta potential measurements of bacteria suspended in MB (n = 3) measure the surface charge of the cells. (C) Comparison bar graph of *Δ2nL* (%) PRISM values for OX and AMINE after 120 min of accumulation time. (D) Individual real-time PRISM accumulation curves for *E. coli* WT, *E. coli* K-12, and *S. epidermidis* on AMINE (positively charged) pore and pillar substrates. Note that the scale of the *E. coli* WT attachment curves is 5-times greater than that of *E. coli* K-12 and *S. epidermidis.* (E) False-colored SEM images of (i) *E. coli* WT on AMINE pores. (ii) *E. coli* WT on AMINE pillars reveal bridging between pillars and elongation of *E. coli* cells, in contrast to (iii) *E. coli* K-12 on AMINE pillars and (iv) *S. epidermidis* on AMINE pillars.

While generally all bacteria exhibit a negative cell surface charge,^[54]^ of the bacteria included in our assays, *S. epidermidis* exhibits the most positive charge, while *E. coli* WT has the most negative charge. These differences may arise due to differences in the peptidoglycan layer, the presence and structure of lipopolysaccharides, and/or the presence of appendages such as flagella (insinuating that charge is not homogenous throughout a cell).^[54,55]^ Differences in measured cell charges may also result from inhomogeneity within the cell population depending on the life cycle stage of the cells.^[56]^ Interestingly, even within the same species, *E. coli* WT was statistically more negatively charged than the K-12 strain. These differences in bacterial cell charge can be seen in the real-time adhesion PRISM curves, presented in Figure 4D, as well as in the attained average *Δ2nL* (%) values after 120 min (Figure 3C). *E. coli* WT exhibited a ~2.7 times greater averaged *Δ2nL* (%) values on positively charged pore surfaces than on the negatively charged pore surfaces (Figure 4C), while K-12 yielded an average *Δ2nL* (%) after 120 min that was ~2.6 times greater on oxidized negatively charged pore surfaces versus positively charged pores. *S. epidermidis*, the most positively charged cells, also exhibited more adhesion to OX surfaces (negative) as opposed to AMINE surfaces (positive). It is important to note that the differently charged surfaces only had minor differences in contact angles, as illustrated in Figure 1A.

While a comparison of numbers may not concretely conclude a relationship between cell and surface charge, complimentary electron microscopy studies support these results, as Figure 4E presents micrographs of the different bacterial species on AMINE surfaces. *E. coli* WT cells elongated and stretch from pillar to pillar or wrap around the pore walls (Figure 4Eii and 4Ei, respectively), perhaps as a way to maximize surface coverage or possibly as the beginning of a filamentation stage that may occur during the initial stages of biofilm formation.^[57]^ Furthermore, optical microscopy of *E. coli* WT on amine-functionalized planar substrates (see Supplemental Movie 1) reveals cells spinning in circles while attached to the surface, suggesting that the AMINE surfaces promote tethering of cells to the surface and by particular locations of the cell body. More specifically, the spinning of all the cells in a counterclockwise direction suggests tethering of the flagella to the surface^[58–60]^ and incidentally with the bacterial strain containing the most flagella per cell, namely *E. coli* WT (as observed in Figure 3C).

### 2.5. Cell-to-cell communication promotes cohesion of bacterial communities

After bacterial cells colonize a surface, often pathogenic strains will then proceed to form intertwined networks of communities in the process of biofilm formation. Thus, we lastly investigate the role of cell-to-cell adhesion in the accumulation of cells on a substrate. For these studies, a variety of pathogenic bacterial strains were employed, including *S. aureus, Enterococcus faecalis* (*E. faecalis*), *Klebsiella pneumoniae* (*K. pneumoniae*), and *Pseudomonas aeruginosa* (*P. aeruginosa*). While all of these strains are clinical isolates that have been maintained in a lab environment, it is interesting to note that the *P. aeruginosa* was isolated from an intravenous coronary stent of a patient at a neighboring medical center (Bnai Zion Medical Center, Israel). The attachment behaviors of these four clinically relevant isolates are depicted in **Figure 5A** as a heat map in which more saturated colors represented more bacterial attachment after 180 min of accumulation in MB. However, unlike the non-pathogenic bacterial strains, in the testing of the clinical isolates, negative values of *Δ2nL* (%) are surprisingly observed for some of the pathogenic bacteria, particularly in the case of *P. aeruginosa* to AMINE pores.

**Figure 5.**
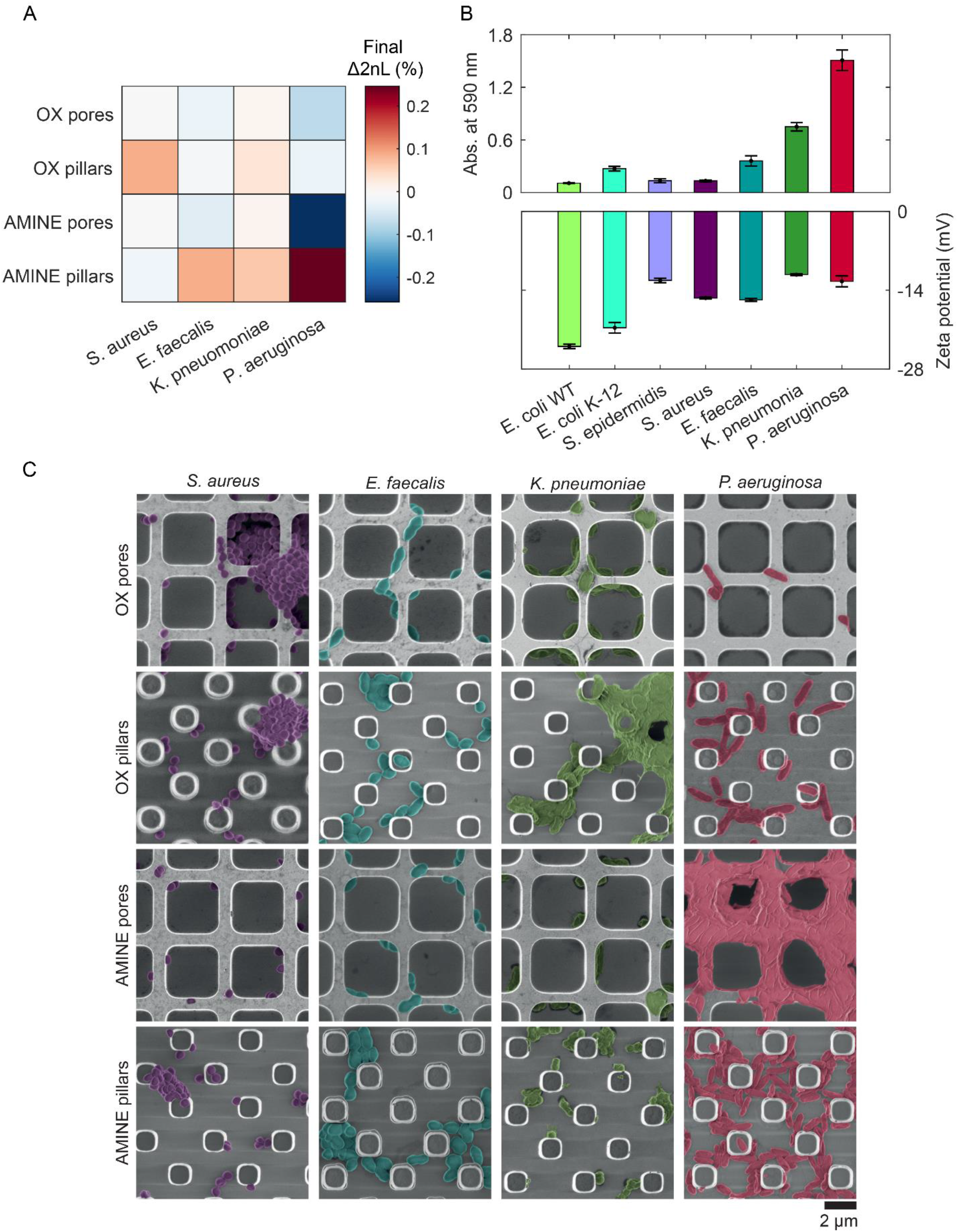
Investigation of pathogenic, clinically relevant bacterial strains (*S. aureus, E. faecalis, K. pneumoniae*, and *P. aeruginosa*). (A) Heat map representing the attained *Δ2nL* (%) values after 180 min of the PRISM accumulation curves for *S. aureus, E. faecalis, K. pneumoniae*, and *P. aeruginosa* on oxidized (OX) and amine-functionalized (AMINE) pore and pillar microtopologies. More saturated colors correspond to denser populated topologies. (B) Biofilm formation abilities of bacteria are quantified by absorbance measurements at 590 nm after staining with crystal violet (top graph). Higher absorption values indicate the presence of more exopolysaccharides and thus, a greater degree of biofilm formation. The zeta potential measurements of the bacterial strains (bottom graph) reveal different cell surface charges for different species. (C) Cohesion of bacterial cells to each other and the substrate after 180 min can also be observed in SEM images (false-colored for clarity).

To understand this behavior of these bacterial strains in-depth, cell surface zeta potentials are measured (n = 3), see Figure 5B. Unlike the *E. coli* strains, cell surface charge did not directly correlate with accumulation on oppositely charged surfaces as measured by PRISM, suggesting that surface heterogeneity on individual cells and also phenotypic heterogeneity within populations may play an even greater role in pathogenic bacteria.^[30,56,61,62]^ However, using a standard crystal violet biofilm formation assay,^[63,64]^ we were able to measure the ability of bacterial strains to develop biofilms by the staining of secreted exopolysaccharides and cells adhered to the bottom of a 96-well plate after the removal of planktonic cells (Figure 5B). As expected, the clinically isolated, highly pathogenic *P. aeruginosa* exhibited the highest biofilm forming abilities as seen in Figure 5C, which corresponds well to the extreme PRISM values (Figure 4A). Furthermore, SEM images of the substrates during the PRISM assay after 180 min of cell accumulation reveal interesting interactions of both cells with each other and cells with the substrate. For example, *S. aureus* forms clusters of cells, while *E. faecalis* arranges into chains of cells. *K. pneumoniae* appears to develop multiple phenotypes with excreted substances, including some cells with numerous fimbriae. *P. aeruginosa* also acquires multiple phenotypes of cells, as well as the initiation of biofilm formation, during which it is appears as if the cells migrate from inside to the top of the pore walls. Such changes in phenotype are known to occur when cells contact a surface.^[1]^ Additionally, the formation of the biofilm on top of the AMINE pores, as opposed to inside the pores, suggests that the excessive presence of cells attached outside the pores can lead to negative values of *Δ2nL* (%) that are only observed with the biofilm-forming strains. The precise mathematical reasoning for this decrease in terms of physics and optical path differences remains unknown. Interestingly, from the observation of these bacteria, we were able to conclude that cells that work together in biofilm formation tend to form the largest, thriving communities (e.g., *P. aeruginosa*) on Si surfaces and that PRISM can be used to predict their biofilm forming abilities within 3 hours. This is significantly faster than the current methods used, such as direct colony forming units counting, crystal violet staining, mass measurements, and flow cytometry of fully formed biofilms, and does not require labeling as does microscopy often does. Additionally, employing PRISM as a method for early detection of biofilm formation would also prove useful for studying the susceptibility of biofilms to antimicrobials/antibiotics.^[65]^

### 2.6. Addressing statistical significance

The statistical significance between all PRISM assays performed using *E. coli* WT, *E. coli* K-12, *S. epidermidis*, and *E. coli* mutants, are summarized in the form of p-values derived from multiway analysis of variance (ANOVA) for testing the effects of surface chemistry, topology, and bacterial species on the mean value of *Δ2nL* (%) acquired after 120 minutes (Figure S2). However, these results should be interpreted with caution.^[66]^ While some of the comparisons reveal that there are differences with p-values < 0.05, such as *E. coli* WT versus non-flagellated E. *coli* on oxidized pores (suggesting that motility may be crucial for attachment on surfaces with limited wetting) we believe that categorizing statistically significant relationships in this study would be misleading, as heterogeneity in bacterial populations is what drives their evolution. Additionally, some conditions with high variances suggest that something in that conditions may bring about more variation than other conditions (*i.e., E. coli* WT on amine-functionalized pillars). Because bacteria cells are so responsive to their environment, growth conditions, and to the status of their neighboring cells, comparisons of p-values with the standard threshold of 0.05 can prove challenging in studies of reproducible biofilms and bacterial attachment.

## 3. Conclusion

The intricacies of bacterial colonization and biofilm formation have become a topic of high importance for medical fields and microbial ecologists. However, finding methods and platforms to do so in a quantitative and label-free way still remains challenging. Thus, we devised a unique platform for real-time monitoring of bacterial attachment in a controlled environment without the use of fluorescent labels or imaging. This platform is based on periodic arrays of microstructures, resulting in photonic diffractive gratings. With these materials, an extensive study of bacteria behavior can be performed.

We found that substrate wetting, which is dependent of surface topology, can critically affect the potential of cells to reach a surface. Cell motility will guide cells to the surface if the wetting of the substrate permits. Furthermore, the charge of the surface can play a role in cell attachment once the cells have reached the surface. However, cell surface charge is not homogenous and may not be homogenous within populations, leading to a more complex cell-surface interfaces than opposite charges will attract. Lastly, the cooperation of cells working as a team to develop complex communities will have a direct effect on the accumulation of cells on a microstructured surface. While many studies have focused on anti-biofouling materials based on observations of non-pathogenic materials, these studies demonstrate that pathogenic bacteria can have extremely different responses. While colloidal theories can compare cells to particles, these theories overlook the complex networks the cells development as communities in order to thrive under various conditions.

While we focus on factors such as surface charge, substrate topology, and substrate wetting, there are still infinite other factors that can be studied with these materials and platform. Additional mechanical factors, such as the possession of bacterial fimbriae, and the role of the surrounding fluid (e.g. temperature, ionic strength, pH, and flow) are a few other factors that can be extensively monitored with this platform. However, we demonstrate that both the nature of the bacteria themselves and the characteristics of the environment around them, specifically the abiotic surface, is what results (or dissuades) bacteria from sticking and forming larger communities on the material. And by simple modifications of materials, we have demonstrated that the maturation of cells and development of bacterial communities is the combined result of the bacteria cells’ nature and substrates’ nurturing environment.

## 4. Experimental Section

### Reagents and solutions

Phenol, tannic acid, crystal violet, sodium lactacte, L-methionine, ethylenediaminetetraacetic acid (EDTA), 3-(aminopropyl)triethoxysilane (APTES), glutaraldehyde, and tryptic soy broth (TSB) were purchased from Sigma-Aldrich, Israel. Acetone and methanol were supplied by Gadot, Israel. Agar was supplied by Difco. Aluminum potassium sulfate dodecahydrate was supplied by Scharlau, Spain. All motility buffer (MB) salts and absolute ethanol were supplied by Merck, Germany. Acetic acid was supplied by Bio-Lab Ltd, Israel. All aqueous solutions were prepared in Milli-Q water (18.2 MΩ cm). Motility buffer (MB) consisted of 6.2 mM K_2_HPO_4_, 3.8 mM KH_2_PO_4_, 67 mM NaCl, 0.1 mM EDTA, 1 μM L-methionine, and 10 mM sodium lactate.

### Fabrication of photonic chips

Si photonic chips containing either lamellar arrays of micropillars or pores were fabricated used standard photolithography and reactive ion etching techniques at the Wolfson Microelectronics Center (Micro- Nano- Fabrication and Printing Unit, Technion – Israel Institute of Technology). Developed whole wafers containing pores were thermally oxidized at 850 °C for 21 min in a tube diffusion furnace (BDF-4 RTRI-878, Bruce Technologies Inc, USA). Each developed wafer was then coated with photoresist to protect the microstructures and diced into 4 mm x 4 mm chips by an automated dicing saw (DAD3350; Disco, Japan). Chips were cleaned in acetone. Wafers containing micropillars were diced, followed by thermal oxidation in a tube furnace (Lindberg/Blue M 1200°C Split-Hinge; Thermo Scientific, USA) at 800 °C for 1 h in ambient air. The resulting oxidized samples are termed “OX.” Amine-modified chips (termed “AMINE”) were generated by incubation of the oxidized samples in 2% v/v APTES in 50% methanol for 1 h, after which they were washed with ethanol and dried under nitrogen.

### Bacterial strains and growth conditions

Five different bacterial species were evaluated, including *Escherichia coli* (including the strains *E. coli* ATCC 25922, K-12, HCB437, HCB1394, and HCB137), *Staphylococcus aureus* ATCC 25923, *Staphylococcus epidermidis* ATCC 14990, *Klebsiella pneumoniae* ATCC 700603, *Enterococcus faecalis* ATCC 29212, and *Pseudomonas aerugonisa* (clinical isolate collected by Bnai Zion Medical Center; Haifa, Israel). All ATCC bacterial strains were obtained from American Type Culture Collection, USA. HCB strains were kindly donated by Dr. Karen Fahrner and Prof. Howard Berg (Harvard University; Cambridge, MA). *E. coli* K-12 was generously donated by Prof. Sima Yaron (Technion – Israel Institute of Technology). All cultures were maintained on tryptic soy agar (TSA) at 4 °C. Each bacterial strain was cultured individually in tryptic soy broth (TSB) overnight at 37 °C, followed by sub-culturing in a 1:100 dilution until an optical density measured at a wavelength of 600 nm (OD_600_) value of ~0.5. All HCB strains were cultured at 30 °C.

### Adhesion assays using PRISM

Bacterial cultures were centrifuged at 2575 g for 5 min at room temperature. The resulting pellet was resuspended in filtered MB and adjusted to a concentration of 10^7^ cells mL^−1^ as determined by calculated OD600 curves. For each adhesion trial, one PRISM chip was fixed in a polycarbonate flow cell at room temperature. A broadband light source (LS1, Ocean Optics) connected to a bifurcated fiber fitter with a collimated lens was focused normal to the surface of the chip. Reflectance spectra were collected by a USB4000 spectrometer (Ocean Optics) every 2 min. Bacterial suspensions were introduced into the flow chamber at a speed of 20 μL min^−1^. PRISM was expressed as *Δ2nL* (%) and calculated as follows:

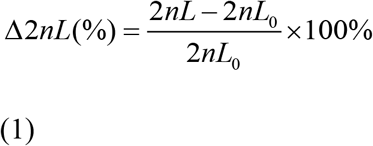

In which *2nL_0_* is the value of *2nL* when the pumping solution is switched to bacterial suspension composed in motility buffer (*t* = 0 of the accumulation time). For each assay, 70% ethanol was briefly injected into the flow cell, followed by continuous flow of MB until stable values of *2nL* were achieved, after which the solution was changed to the bacterial suspension.

### Assays of pathogenic bacteria

Due to the pathogenicity of the strains and desire to observe faster differences in adhesion preferences, assays of pathogenic bacteria, namely *E. faecalis, S. aureus*, *P. aeruginosa*, and *K. pneumoniae*, were modified. In these assays, bacteria were plated on TSA overnight. Colonies were selected from freshly incubated agar plates and suspended in MB. The suspension was diluted until achieving and OD_600_ of 0.1.

### Crystal violet biofilm formation assay

Aliquots of 10 μL of overnight bacterial cultures in TSB were seeded in a 96-well polystyrene plate. Cells were overlaid with 190 μL of TSB and incubated for 24 h at room temperature. The plate was submerged in ddH2O for 5 min to remove planktonic cells, followed by inversion of the plate to remove remaining water drops. Wells were incubated for 10 min with 0.1% crystal violet solution, after which the plate was submerged in diH2O for 5 min to remove excess crystal violet. The plate was again inverted to remove large water droplets containing crystal violet. After drying for 20 min, 200 μL of 96% ethanol was added to each well and incubated for 5 min to solubilized crystal violet contained within the biofilms. 180 μL of the solubilized crystal violet was transferred to a new 96-well plate, where absorbance at 590 nm was measured by a microplate reader (Synergy HT, BioTek Instruments). The average absorbance value of wells inoculated with only growth medium was subtracted from the absorbance values of bacteria-containing wells to calibrate for any excess crystal violet adhered to the well plate.

### Motility assay in semi-solid agar

Semi-solid agar was prepared by adding 0.25% agar to TSB. Following autoclaving, ~20 mL of agar was poured into each plate. Agar plates remained at room temperature overnight. A bacterial suspension of 10 μL was spotted in the middle of each plate and incubated at room temperature to mimic conditions of the optical experiments. Agar plates were imaged and the diameter of each radial growth area was measured using ImageJ software.

### Contact angle measurements

Contact angles of 5 μL drops of MB on various substrates were measured with an Attension Theta Lite optical tensiometer (Biolin Scientific). A CCD camera acquired digital images of the droplet on each substrate over time. Contact angles were calculated by OneAttension Software after 200 s using the Young–Laplace equation. Single measurements were performed on three different chips of the same type as the droplet covered the majority of the substrate surface.

### Zeta potential measurements

Bacterial suspensions taken from freshly incubated plates were prepared to an OD_600_ 0.1 in MB (pH= 6.9). Zeta potential measurements of the suspensions were performed using a ZetaSizer ZSP (Malvern Instruments, UK) in triplicate samples.

### Imaging

All PRISM chips were fixed and stored at 4 °C in 2.5% glutaraldehyde. For scanning electron microscopy (SEM), samples were washed in subsequent 50-70-100% ethanolic solutions, each for 5 min and then dried under a stream of nitrogen. Samples were analyzed at 1 keV using a high-resolution SEM (Ultra Zeiss Plus). To visualize flagella, a Ryu flagellar stain was performed.^23,24^ Briefly, bacterial cells were grown overnight on tryptic soy agar. Drops of motile cells were prepared in sterile water by touching a loopful of water to the colony margin on the agar and then suspending the bacteria by touching a drop of water on a glass slide, which was covered with a cover slip and examined for motile cells. Then, two drops of Ryu stain were applied to the edge of the cover slip and allowed to diffuse by capillary action. The Ryu stain is composed of a filtered 1:10 mixture of two solutions, in which the first solution contains 5% phenol and 20% tannic acid in a saturated aqueous solution of aluminum potassium sulfate dodecahydrate and the second solution contains 12% crystal violet in 95% ethanolic solution. Bacterial cells were examined for flagella after 10 min using a Zeiss Axio Scope A1 microscope. Images were obtained by an Axio Cam MRc (Zeiss) camera.

## Acknowledgements

This work was funded by the Israeli Ministry of Science. We thank Dr. Karen Fahrner and Prof. Howard Berg of Harvard University for donating *E. coli* mutant strains and for their useful discussions and insight. We thank Dima Peselev, Orna Ternyak, and the rest of the staff at the Micro- Nano- Fabrication and Printing Unit (MNFPU) at the Technion – Israel Institute of Technology for fabrication of the diffraction gratings. We thank Prof. Boaz Pokroy and Beni Rich for guidance with contact angle measurements.

