## Supporting Information for "Sticky Situations: Bacterial Attachment Deciphered by Interferometry of Silicon Microstructures"

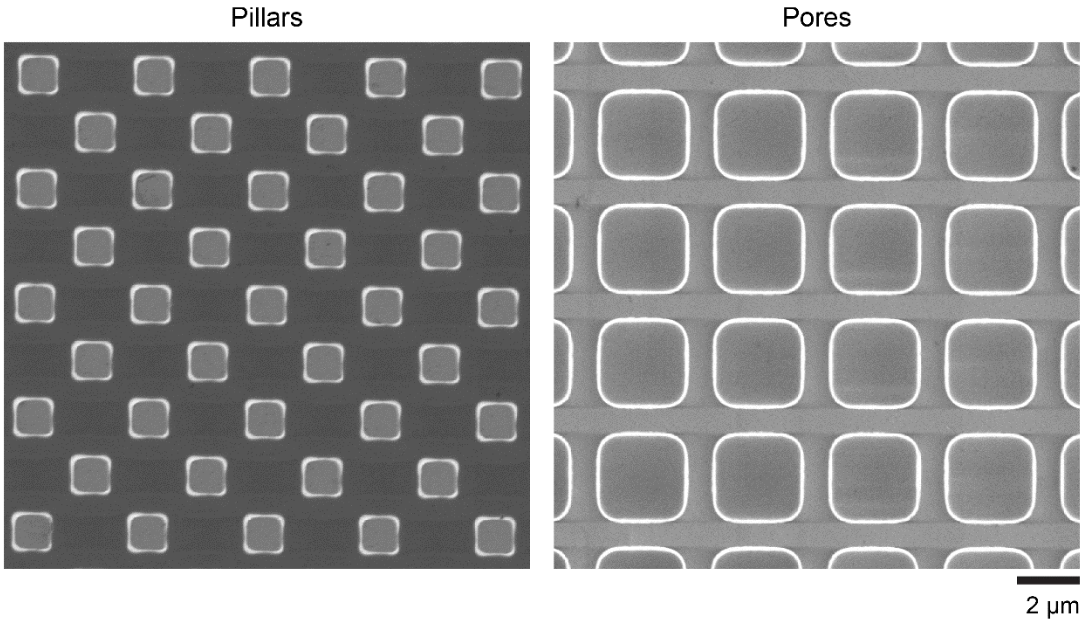


**Figure S1.** Plan-view SEM images of Si pillar microstructures (left) and pore microstructures (right).

**Figure S2.** (Following page) p-values for multiway analysis of variance (ANOVA) for testing the effects of surface chemistry, topology, and bacterial species on the mean value of *Δ2nL* (%) acquired after 120 minutes. OX = oxidized; AMINE = Amine-functionalized; WT = *E. coli* WT; K12 = *E. coli* K-12; SE = *S. epidermidis*; tumbly, no flagella, no chemotaxis refer to mutant *E. coli* strains. Bluer values correspond to comparisons of mean values the lead to lower p-values and are thus more statistically significant.

| | AMINE pillar, WT | AMINE pore, WT | OX pillar, WT | OX pore, WT | AMINE pillar, K12 | AMINE pore, K12 | OX pillar, K12 | OX pore, K12 | AMINE pillar, SE | AMINE pore,SE | OX pillar,SE | OX pore, SE | AMINE pillar, tumbly | AMINE pore, tumbly | OX pillar, tumbly | OX pore, tumbly | AMINE pillar, no flagella | AMINE pore, no flagella | OX pillar, no flagella | OX pore, no flagella | AMINE pillar, no chemotaxis | AMINE pore, no chemotaxis | OX pillar, no chemotaxis | OX pore,no chemotaxis |
|---|---|---|---|---|---|---|---|---|---|---|---|---|---|---|---|---|---|---|---|---|---|---|---|---|
| AMINE pillar, WT | | 0.000 | 0.099 | 1.000 | 0.215 | 0.120 | 0.027 | 0.558 | 0.256 | 0.182 | 0.907 | 0.036 | 0.042 | 0.052 | 0.064 | 0.054 | 0.019 | 0.115 | 0.054 | 0.023 | 0.034 | 0.025 | 0.103 | 1.000 |
| AMINE pore, WT | 0.000 | | 0.000 | 0.000 | 0.000 | 0.000 | 0.000 | 0.000 | 0.000 | 0.000 | 0.000 | 0.000 | 0.000 | 0.000 | 0.000 | 0.000 | 0.000 | 0.000 | 0.000 | 0.000 | 0.000 | 0.000 | 0.000 | 0.000 |
| OX pillar, WT | 0.099 | 0.000 | | 0.185 | 1.000 | 1.000 | 1.000 | 1.000 | 1.000 | 1.000 | 0.997 | 1.000 | 1.000 | 1.000 | 1.000 | 1.000 | 1.000 | 1.000 | 1.000 | 1.000 | 1.000 | 1.000 | 1.000 | 0.206 |
| OX pore, WT | 1.000 | 0.000 | 0.185 | | 0.360 | 0.219 | 0.057 | 0.746 | 0.415 | 0.313 | 0.976 | 0.074 | 0.085 | 0.104 | 0.124 | 0.107 | 0.040 | 0.211 | 0.106 | 0.049 | 0.071 | 0.052 | 0.192 | 1.000 |
| AMINE pillar, K12 | 0.215 | 0.000 | 1.000 | 0.360 | | 1.000 | 1.000 | 1.000 | 1.000 | 1.000 | 1.000 | 1.000 | 1.000 | 1.000 | 1.000 | 1.000 | 1.000 | 1.000 | 1.000 | 1.000 | 1.000 | 1.000 | 1.000 | 0.392 |
| AMINE pore, K12 | 0.120 | 0.000 | 1.000 | 0.219 | 1.000 | | 1.000 | 1.000 | 1.000 | 1.000 | 0.998 | 1.000 | 1.000 | 1.000 | 1.000 | 1.000 | 1.000 | 1.000 | 1.000 | 1.000 | 1.000 | 1.000 | 1.000 | 0.242 |
| OX pillar, K12 | 0.027 | 0.000 | 1.000 | 0.057 | 1.000 | 1.000 | | 0.999 | 1.000 | 1.000 | 0.928 | 1.000 | 1.000 | 1.000 | 1.000 | 1.000 | 1.000 | 1.000 | 1.000 | 1.000 | 1.000 | 1.000 | 1.000 | 0.065 |
| OX pore, K12 | 0.558 | 0.000 | 1.000 | 0.746 | 1.000 | 1.000 | 0.999 | | 1.000 | 1.000 | 1.000 | 1.000 | 1.000 | 1.000 | 1.000 | 1.000 | 0.995 | 1.000 | 1.000 | 0.998 | 0.999 | 0.998 | 1.000 | 0.777 |
| AMINE pillar, SE | 0.256 | 0.000 | 1.000 | 0.415 | 1.000 | 1.000 | 1.000 | 1.000 | | 1.000 | 1.000 | 1.000 | 1.000 | 1.000 | 1.000 | 1.000 | 1.000 | 1.000 | 1.000 | 1.000 | 1.000 | 1.000 | 1.000 | 0.448 |
| AMINE pore,SE | 0.182 | 0.000 | 1.000 | 0.313 | 1.000 | 1.000 | 1.000 | 1.000 | 1.000 | | 1.000 | 1.000 | 1.000 | 1.000 | 1.000 | 1.000 | 1.000 | 1.000 | 1.000 | 1.000 | 1.000 | 1.000 | 1.000 | 0.342 |
| OX pillar,SE | 0.907 | 0.000 | 0.997 | 0.976 | 1.000 | 0.998 | 0.928 | 1.000 | 1.000 | 1.000 | | 0.954 | 0.966 | 0.979 | 0.987 | 0.981 | 0.876 | 0.998 | 0.980 | 0.906 | 0.950 | 0.915 | 0.997 | 0.982 |
| OX pore, SE | 0.036 | 0.000 | 1.000 | 0.074 | 1.000 | 1.000 | 1.000 | 1.000 | 1.000 | 1.000 | 0.954 | | 1.000 | 1.000 | 1.000 | 1.000 | 1.000 | 1.000 | 1.000 | 1.000 | 1.000 | 1.000 | 1.000 | 0.083 |
| AMINE pillar, tumbly | 0.042 | 0.000 | 1.000 | 0.085 | 1.000 | 1.000 | 1.000 | 1.000 | 1.000 | 1.000 | 0.966 | 1.000 | | 1.000 | 1.000 | 1.000 | 1.000 | 1.000 | 1.000 | 1.000 | 1.000 | 1.000 | 1.000 | 0.096 |
| AMINE pore, tumbly | 0.052 | 0.000 | 1.000 | 0.104 | 1.000 | 1.000 | 1.000 | 1.000 | 1.000 | 1.000 | 0.979 | 1.000 | 1.000 | | 1.000 | 1.000 | 1.000 | 1.000 | 1.000 | 1.000 | 1.000 | 1.000 | 1.000 | 0.117 |
| OX pillar, tumbly | 0.064 | 0.000 | 1.000 | 0.124 | 1.000 | 1.000 | 1.000 | 1.000 | 1.000 | 1.000 | 0.987 | 1.000 | 1.000 | 1.000 | | 1.000 | 1.000 | 1.000 | 1.000 | 1.000 | 1.000 | 1.000 | 1.000 | 0.140 |
| OX pore, tumbly | 0.054 | 0.000 | 1.000 | 0.107 | 1.000 | 1.000 | 1.000 | 1.000 | 1.000 | 1.000 | 0.981 | 1.000 | 1.000 | 1.000 | 1.000 | | 1.000 | 1.000 | 1.000 | 1.000 | 1.000 | 1.000 | 1.000 | 0.121 |
| AMINE pillar, no flagella | 0.019 | 0.000 | 1.000 | 0.040 | 1.000 | 1.000 | 1.000 | 0.995 | 1.000 | 1.000 | 0.876 | 1.000 | 1.000 | 1.000 | 1.000 | 1.000 | | 1.000 | 1.000 | 1.000 | 1.000 | 1.000 | 1.000 | 0.046 |
| AMINE pore, no flagella | 0.115 | 0.000 | 1.000 | 0.211 | 1.000 | 1.000 | 1.000 | 1.000 | 1.000 | 1.000 | 0.998 | 1.000 | 1.000 | 1.000 | 1.000 | 1.000 | 1.000 | | 1.000 | 1.000 | 1.000 | 1.000 | 1.000 | 0.234 |
| OX pillar, no flagella | 0.054 | 0.000 | 1.000 | 0.106 | 1.000 | 1.000 | 1.000 | 1.000 | 1.000 | 1.000 | 0.980 | 1.000 | 1.000 | 1.000 | 1.000 | 1.000 | 1.000 | 1.000 | | 1.000 | 1.000 | 1.000 | 1.000 | 0.120 |
| OX pore, no flagella | 0.023 | 0.000 | 1.000 | 0.049 | 1.000 | 1.000 | 1.000 | 0.998 | 1.000 | 1.000 | 0.906 | 1.000 | 1.000 | 1.000 | 1.000 | 1.000 | 1.000 | 1.000 | 1.000 | | 1.000 | 1.000 | 1.000 | 0.055 |
| AMINE pillar, no chemotaxis | 0.034 | 0.000 | 1.000 | 0.071 | 1.000 | 1.000 | 1.000 | 0.999 | 1.000 | 1.000 | 0.950 | 1.000 | 1.000 | 1.000 | 1.000 | 1.000 | 1.000 | 1.000 | 1.000 | 1.000 | | 1.000 | 1.000 | 0.080 |
| AMINE pore, no chemotaxis | 0.025 | 0.000 | 1.000 | 0.052 | 1.000 | 1.000 | 1.000 | 0.998 | 1.000 | 1.000 | 0.915 | 1.000 | 1.000 | 1.000 | 1.000 | 1.000 | 1.000 | 1.000 | 1.000 | 1.000 | 1.000 | | 1.000 | 0.059 |
| OX pillar, no chemotaxis | 0.103 | 0.000 | 1.000 | 0.192 | 1.000 | 1.000 | 1.000 | 1.000 | 1.000 | 1.000 | 0.997 | 1.000 | 1.000 | 1.000 | 1.000 | 1.000 | 1.000 | 1.000 | 1.000 | 1.000 | 1.000 | 1.000 | | 0.213 |
| OX pore,no chemotaxis | 1.000 | 0.000 | 0.206 | 1.000 | 0.392 | 0.242 | 0.065 | 0.777 | 0.448 | 0.342 | 0.982 | 0.083 | 0.096 | 0.117 | 0.140 | 0.121 | 0.046 | 0.234 | 0.120 | 0.055 | 0.080 | 0.059 | 0.213 | |

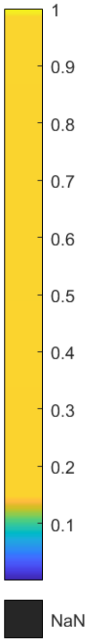
